# Modeling genotype × environment interaction for single- and multi-trait genomic prediction in potato (*Solanum tuberosum* L.)

**DOI:** 10.1101/2022.08.09.503418

**Authors:** Jaime Cuevas, Fredrik Reslow, Jose Crossa, Rodomiro Ortiz

## Abstract

In this study we extend research on genomic prediction (GP) to polysomic polyploid plant species with the main objective to investigate single trait (ST) versus multi-trait (MT) for multi-environment (ME) models for the combination of three locations in Sweden (Helgegården [HEL], Mosslunda [MOS], Umeå [UM]) over two year-trials (2020, 2021) of 253 potato cultivars and breeding clones for five tuber weight traits and two tuber flesh quality characteristics. This research investigated the GP of four genome-based prediction models with genotype ×environment interactions (GE): (1) single trait reaction norm model (*M1*), (2) single trait model considering covariances between environments (*M2*), (3) single trait *M2* extended to include a random vector that utilizes the environmental covariances (*M3*) and (4) multi-trait model with GE (*M4*). Several prediction problems were analyzed for each of the GP accuracy of the four models. Results of the prediction of traits in HEL, the high yield potential testing site in 2021, show that the best predicted traits were tuber flesh starch (%), weight of tuber above 60 or below 40 mm in size, and total tuber weight. In terms of GP, accuracy model *M4* gave the best prediction accuracy in three traits, namely tuber weight of 40–50 or above 60 mm in size, and total tuber weight and very similar in the starch trait. For MOS in 2021, the best predictive traits were starch, weight of tuber above 60, 50–60, or below 40 mm in size, and total tuber weight. MT model *M4* was the best GP model based on its accuracy when some cultivars are observed in some traits. For GP accuracy of traits in UM in 2021, the best predictive traits were weight of tuber above 60, 50–60, or below 40 mm in size and the best model was MT *M4* followed by models ST *M3* and *M2*.

## 1. INTRODUCTION

Genomic prediction (GP) and selection (GS) have changed the paradigm of plant and animal breeding (Meuwissen et al., 2001; de los Campos et al., 2009; Crossa et al., 2010, 2011, Desta and Ortiz, 2014). Practical evidence has shown that GS provides important increases in prediction accuracy for genomic-aided breeding (Crossa et al., 2014, 2017; Pérez-Rodríguez et al., 2012). Additive genetic effects (breeding values) can be predicted directly from parametric and semi-parametric statistical models using marker effects like the ridge regression best linear unbiased prediction (rrBLUP) (Endelman, 2011), or by developing the genomic relationship linear kernel matrix (***G***) to fit the genomic best linear unbiased prediction [GBLUP] (VanRaden, 2008). Departures from linearity can be assessed by semi-parametric approaches, such as Reproducing Kernel Hilbert Space (RKHS) regression using the Gaussian kernel or different types of neural networks (Gianola et al., 2006; Gianola and van Kaam, 2008; de los Campos et al., 2010; González-Camacho et al., 2012; Pérez-Rodríguez et al., 2012, Gianola et al., 2014; Sousa et al., 2017).

Standard GP models were extended to multi-environments by assessing genomic × environment interaction (GE) (Burgueño et al., 2012). Jarquín et al. (2014) proposed an extension of the GBLUP or random effects model where the main effects of markers and environmental covariates could be introduced using covariance structures that are functions of marker genotypes and environments. Consistently, GP accuracy substantially increased when incorporating GE and marker × environment interaction (Crossa et al., 2017). Cuevas et al. (2016) and Souza et al. (2017) applied the marker × environment interaction GS model of Lopez-Cruz et al. (2015) but modeled not only through the standard GBLUP but also through a non-linear Gaussian kernel (GK) like that used by de los Campos et al. (2010) and a GK with the bandwidth estimated through an empirical Bayesian method (Pérez-Elizalde et al., 2015). Cuevas et al. (2016) concluded that the higher prediction accuracy of the GK models with the GE model is due to more flexible kernels that allow accounting for small, more complex marker main effects and marker-specific interaction effects.

In GP the training set usually includes a sufficient overlap of lines across environments, so that estimating the phenotypic covariance among environments for modeling GE is sufficient to specify it on the linear mixed model used. When modeling GE, some researchers used the mathematical operation defined by the Kronecker products or direct product (Cuevas et al., 2016) that allows operations of two matrices of different dimensions. Other authors model GE using the matrix operation named Hadamard products (also known as element-wise products) that is a binary operation between two matrices of the same dimensions as the operands (Jarquin et al., 2014; Lopez-Cruz et al., 2015; Perez-Rodriguez et al., 2015; Acosta-Pech et al., 2017; Perez-Rodriguez et al., 2017; Sukumaran et al. 2017; Basnet et al., 2019). When modeling epistasis, Hadamard products of the additive genomic relationship have mainly been used (e.g., Jiang and Reif, 2015; Martini et al., 2016; Vitezica et al., 2017; Varona et al., 2018; Martini et al., 2020). However, Crossa et al. (2006) and Burgueño et al. (2007) have used Kronecker products for modeling and estimation of additive, additive × environment interaction, additive × additive epistasis, and additive × additive × environment interactions by means of the coefficient of parentage. In a recent study, Martini et al. (2020) gave theoretical proof that both methods lead to the same covariance model when used with some specific design matrices and illustrated how to explicitly model the interaction between markers, temperature, or precipitation.

Traditionally GP models have evolved from the single trait (ST) and single environment prediction (ST-SE) models to ST multiple environment (ST-ME) models including GE. Furthermore, standard GS-assisted plant breeding models are concerned with the assessment of the GP accuracy of a multi-trait (MT) measured in a single environment (MT-SE) or MT multiple environments (MT-ME). In general, multi-traits (MT) GP models have evolved from MT-SE to MT-ME. The MT models are key for improving prediction accuracy in GS because MT models offer benefits regarding ST models when the traits under study are correlated. Most existing models for genomic prediction are ST models although MT models have several advantages over the ST (Montesinos et al., 2019). Compared with ST, MT can simultaneously exploit the correlation between cultivar and traits and thus improve the accuracy of GP as they are computationally more efficient than ST (Montesinos-López et al., 2019). When the traits are correlated, MT models improve parameter estimates and prediction accuracy as compared to ST models (Schulthess et al. 2018; Calus and Veerkamp, 2011, Jiang and Jannink, 2012, Montesinos-López et al., 2016, 2019; He et al., 2016). With the continuous growth of computational power, MT models play an increasingly important role in data analysis in plant and animal genomic−aided breeding for selecting the best candidate genotypes.

The use of MT models is not as widespread as the use of ST models because several factors such as, among others, lack of efficient and friendly software, and not enough computational resources; also, MT models have more complex genotype × environment interactions (GE) that make it difficult to assess and achieve MT model assumptions, and MT models have more problems of convergence than ST models. Some models have been proposed for MT GP, e.g., multi-trait mixed models and their Bayesian version, Bayesian multi-trait genomic best linear unbiased predictor and multi-trait models under artificial deep neural networks applied to maize and wheat datasets (Montesinos-López et al., 2018, 2019). However, most researchers use MT models to improve prediction accuracy for traits to be predicted (i.e., the prediction set) −which are tedious and time-consuming to measure and have low heritability− by using a few traits (i.e., the training set) with high heritability that are highly correlated with the former prediction set (Semagn et al., 2022; Jiang and Jannink, 2012).

It is widely recognized that from the statistical and quantitative genetics perspectives, when data on multi-traits are available, the preferred models are the MT as they can account for correlations between phenotypic traits in the training set because borrowing information from correlated traits increases GP accuracy. Montesinos et al. (2022) investigated Bayesian multi-trait kernel methods for GP and illustrated the power of linear, Gaussian, polynomial, and sigmoid kernels. The authors compared these kernels with the conventional ridge regression and GBLUP multi-trait models. Montesinos et al. (2022) showed that, in general, but not always, the GK method outperformed conventional Bayesian ridge and GBLUP multi-trait in terms of GP prediction performance; the authors concluded that the improvement in terms of prediction performance of the Bayesian multi-trait kernel method can be attributed to the proposed model being able to capture nonlinear patterns more efficiently than linear multi-trait models.

Semagn et al. (2022) were interested in comparing prediction accuracy estimates of a subset of lines that have been tested for a single trait (ST), with a subset of lines that have not been tested for certain proportion traits (MT1, certain cultivars were not tested for any of the traits), and a subset of lines that have been tested for some traits but not for other traits (MT2) across different bread wheat genetic backgrounds for agronomic traits of varying genetic architecture evaluated under conventional and organic management systems, and several host plant resistance traits evaluated in adult plants under standard field management. Their results show that the predictive ability of the MT2 model was significantly greater than that of the ST and MT1 models for most of the traits and populations, except common bunt, with the MT1 model being intermediate between them, demonstrating the high potential of the multi-trait models in improving prediction accuracy.

Although most GP research for ST and MT for SE or ME has been applied to diploid species, a recent study by Ortiz et al. (2022) demonstrated the increase in prediction accuracy of ST-ME over the ST-SE genomic-estimated breeding values for several tetrasomic potato (*Solanum tuberosum* L.) breeding clones and released cultivars for various traits evaluated in several sites for one year. Ortiz et al. (2022) considered four dosages of marker alleles (A) pseudo-diploid; (B) additive tetrasomic polyploidy, and (C) additive-non-additive tetrasomic polyploidy, and B+C dosages together in the genome-based prediction models using the conventional linear GBLUP (GB) and the non-linear Gaussian kernel (GK) for ST-SE and ST-ME together. Results show that GK did not show any clear advantage over GB, and ST-ME had prediction accuracy estimates higher than those obtained from ST-SE. The model with GB was the best method in combination with the marker structures C or B+C for predicting most of the tuber traits. Most of the traits gave relatively high prediction accuracy under this combination of marker structure C or (B+C) and methods GB and GK combined with the ST-ME including GE model.

Based on the above considerations, and the need to extend research on GP to polysomic polyploid plant species, the main objectives of this research were to investigate ST versus MT for ME (GE) models for the combination of three locations (namely Helgegården [HEL], Mosslunda [MOS], and Umeå [UM]) over two years (2020, 2021) of 253 potato cultivars and breeding clones, which were also included by Ortiz et al. (2022). In this study we will use only the genomic relationship matrix obtained from the additive-non-additive tetrasomic polyploidy (C) because using this genomic relations matrix in terms of GP accuracy was found to be one with the best GP accuracy (Ortiz et al., 2022). This research investigated the GP of four genome-based prediction models including either Hadamard or Kronecker product matrices for assessing GE: (1) the conventional reaction norm model incorporating GE with Hadamard product (Jarquin et al., 2014) (*M1*), (2) GE model considering covariances between environments, similar to the model employed by Burgueño et al. (2012) or the GE with Kronecker product (*M2*), (3) GE model 2 including a random vector that attempts to more efficiently utilize the environmental covariances as in Cuevas et al. (2017) or a GE with Kronecker product (*M3*), and (4) a multi-trait model with GE as in Montesinos et al. (2022) but including a GE model that joins Hadamard and Kronecker products (*M4*). Several prediction problems were analyzed for the GP accuracy of each of the four models. We investigated the prediction set of locations in year 2021 from locations in year 2020 using the four GP models combined with two of the prediction sets (100% and 70%) and predicting ST and MT.

## 2. MATERIALS AND METHODS

### 2.1 Phenotypic data

The multi-site experiments included 253 potato breeding clones and cultivars in trials at Helgegården (HEL), Mosslunda (MOS) and Umeå (UM). Their list is provided by Ortiz et al. (2022) **Supplementary Table S1** (https://hdl.handle.net/11529/10548617). The breeding clones are in at least the fourth generation (T_4_) of selection by Svensk potatisförädling of the Swedish University of Agricultural Sciences (Ortiz et al., 2020), while the cultivars are a sample of those released and grown in Europe during the last 200 years. Helgegården and Mosslunda are near Kristianstad (56°01′46″N 14°09′24″E, Skåne, southern Sweden), while Umeå (63°49′30″N 20°15′50″E) is in the north of Sweden.

An incomplete block design (simple lattice) with two replications of 10 plants each was the field layout for the field trials across testing sites. Fungicides were only used in Helgegården to avoid late blight caused by the oomycete *Phytophthora infestans* throughout the growing season, thus allowing tuber yield potential to be estimated at this site. Crop husbandry was that used for potato farming at each site.

Total tuber yield per plot (kg), tuber weight by size (< 40 mm, 40–50 mm, 50–60 mm, > 60 mm; kg), while tuber flesh starch was measured as percentage based on specific gravity after harvest. Reducing sugars in the tuber flesh after harvest was determined using potato glucose strip tests (Mann et al., 1991). Host plant resistance to late blight was evaluated using the area under the disease progress curve (AUDPC) in Mosslunda.

### 2.2 Genotypic data

After sampling using four leaf punches for each of the 256 breeding clones and cultivars included in the experiments, the materials were sent by AgriTech – Intertek ScanBi Diagnostics (Alnarp, Sweden) to Diversity Array Technology Pty Ltd (ACT, Australia) for targeted genotyping following a genotype-by-sequencing approach (https://www.diversityarrays.com/technology-and-resources/targeted-genotyping/). More than 2000 single nucleotide polymorphisms (SNP) were used for genotyping. They derived mostly from SolCAP SNPs based on chromosome positions and MAF > 1 in germplasm from the Centro Internacional de la Papa (CIP, Lima, Perú) and the USA. According to Selga et al. (2021), such a number of SNPs seems to be enough for researching GEBVs without losing information. Although there were very few missing genotyping data (0.1%), one breeding clone (97) and two cultivars (‘Leyla’ and ‘Red Lady’) were not included further in the analysis because they were lacking enough SNP data.

### 2.3 Computing the genomic relationship matrix

We briefly described the method used for codifying the molecular ***X*** matrix proposed by Slater et al. (2016) and used one of the options used by Ortiz et al. (2022) in the genomic-enabled prediction models.

### 2.4 Full tetrasomic including additive and non-additive effects

For coding matrix ***X*** according to Slater et al. (2016), we considered additive and non-additive effects in a full tetrasomic assuming each genotype has its own effect. In this case, there were five possible effects per SNP marker. Then the genomic relationship between individuals *j, k* was computed as:

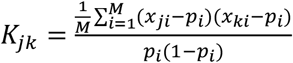

where M was the number of markers × 5. To compute the diagonal of this matrix, we used:

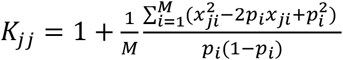

where *p*_*i*_ was the frequency of each genotype, i.e., the frequency in each column.

### 2.5 Statistical models

#### 2.5.1 Single-trait conventional reaction norm model including GE (model 1, *M1*)

The standard reaction norm model incorporating genomic × environment (GE) (Jarquin et al., 2014), as shown below, explains the variation of the observations of a single trait (ST) in each of the *m* environments (ME) represented by the vector ***y*** *=* (***y****′*_*1*_,*…*, ***y****′*_*i*_,*…* ***y****′*_*m*_)′ by estimating each mean of the environment observations ***µ***_*E*_, plus the prediction of the main genetic effects ***g*** and the prediction of the interaction random effects G×E represented by vector ***ge***, the unexplained differences or random errors are represented by vector ***ε***.

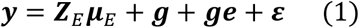

where ***y*** *=* (***y****′*_*1*_,*…*, ***y****′*_*i*_,*…* ***y****′*_*m*_)′ is a column vector of size *n*_*T*_ × *1* of the observations of each environment ***y***_*i*_ (the ’ sign indicates the transpose operation), that is, *n*_*T*_ × *1* is the total of the sum of the number of lines in each environment. The vector ***µ***_*E*_ is a vector that represents the means of the *m* environments, and the incidence matrix ***Z***_*E*_ relates the observations to the mean of the environments. The random genetic vector of main effects ***g*** including GE *n*_*T*_ × 1 follows a multivariate normal distribution 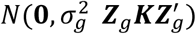 where 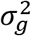 is the variance component of *g*, ***Z***_*g*_ is an incidence matrix that relates the observations with the genotypes and ***K*** is a matrix of relations between the genotypes built with molecular markers. In our study ***K*** was computed as previously indicated for the case of a full tetrasomic genomic relationship matrix. The random vector of interaction effects ***ge*** follows a multivariate normal distribution 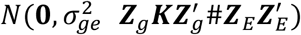 where 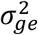 is the variance component and # is the Hadamard product. Random errors are considered with homogeneous variance, that is, 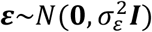. In general, when the correlation between environments in the training set are positive and high, results from using Hadamard product to model GE are similar to those obtained using the Kronecker product. This model is flexible because it allows predicting different numbers of lines in different environments or even predicting the entire environment. However, when the correlations between the environments are not positive, the GE model with the Hadamard product does not explain the phenotype variation well enough (López-Cruz et al., 2015), because the model does not incorporate covariances between environments.

#### 2.5.2 Single trait GE (ST-ME) model considering covariances between environments (model 2, *M2*)

Based on Burgueño et al. (2012), the genomic prediction model including GE considered the genomic covariances between environments to attempt improving the genomic prediction accuracy of unobserved environments. In *M2* we considered only one trait (ST) and multi environments (ME), but the main effect of genomic and the GE interaction effects are modeled jointly by using a single vector ***u*** assuming a multivariate normal distribution that considers the genomics covariances between environments. One form of this model is:

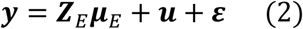

where the vectors ***Z***_*E*_***µ***_*E*_ are similar to those of *M1*, that is, the ***µ***_*E*_ is a vector that represents the means of the *m* environments, and the incidence matrix ***Z***_*E*_ relates the observations with the mean of the environments, but now the number of cultivars is the same for each environment so that if we order the phenotypic observations of the first environment, then the second environment and so forth, 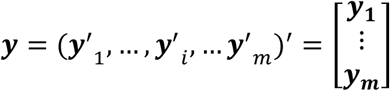; thereafter the genetic random effects can be modeled as a normal distribution ***u****∼N*(**0, *U***_*E*_**⊗*K***), where ***U***_*E*_ is a matrix of genomic covariances between the environments of size *m* × *m* to be estimated, and **⊗** indicates the Kronecker product. The matrix ***K*** represents the relationships between the genotypes built with the molecular markers, as previously indicated. The random errors are modeled as ***ε****∼N*(**0, Σ⊗*I***), where matrix **Σ** is a matrix of size *m* × *m*, expressing the covariances of the errors between environments to be estimated, and ***I*** is the identity matrix of order *n*_*l*_ × *n*_*l*_ (Cuevas et al., 2017). In this study it is assumed that **Σ** is a diagonal matrix that needs to be estimated. Although model *M2* is powerful when considering the genetic covariances between environments, it cannot predict full environments because it does not have a way of estimating the corresponding genomic covariances of those environments in the training sites with those in the testing sites where no data have been collected.

### 2.5.3 Single trait GE model (ST-ME) with an extra random vector to better account for variance across environments (model 3, *M3*)

Cuevas et al. (2017) showed that adding a random vector to *M2* to account for the cultivar variation across environments that was accounted for by vector ***u***, could increase the prediction accuracy. Here we considered a single trait (ST) measured in different environments (ME) to construct and add a random vector ***f*** to *M2*, that is:

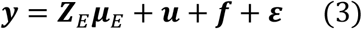

Note that ***y*** is a vector that started with the first environment, then the second environment and so forth until the last environment. Then ***Z***_*E*_***µ***_*E*_ represents the mean for each environment and ***u*** is a random vector with multivariate normal distribution ***u****∼N*(**0, *U***_*E*_**⊗*K***). Then a random vector ***f*** is added that is independent from ***u***, and ***ε***, and that has a normal distribution ***f****∼N*(**0, *F***_*E*_**⊗*I***) where ***F***_*E*_ is a matrix of environmental covariances of size *m* × *m* to be estimated, **⊗** indicates the Kronecker product, and matrix ***I*** represents the identity matrix.

*M3*, like *M2*, allows improving the prediction accuracy of model *M1*, when the covariances (or correlations) of the observations between environments are negative or close to zero. Like *M2, M3* could not be used to predict complete environments because technically it could not estimate covariances between related environments with the environments to be predicted because of the lack of data on the environments to be predicted.

*M2* and *M3* can be used as a multi-trait model for one single site (SE), considering traits instead of environments. In fact, some of the programs for fitting *M2* are motivated by multi-trait models, such as the MTM (multi-trait model) package proposed by de los Campos and Grueneber (2016), and the multi-trait function of the BGLR R-package (de los Campos and Pérez Rodríguez, 2014).

#### 2.5.4 Multi-trait model with GE (model 4, *M4*) of MT-ME type

Note that *M2* could be adopted to be a single environment multi-trait (MT-SE) as

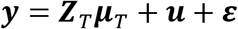

where the vectors ***Z***_*T*_***µ***_*T*_ are similar to those of M2, that is, the ***µ***_*T*_ is a vector that represents the means of the *t* traits, and the incidence matrix ***Z***_*T*_ relates the observations with the mean of the traits, but now the number of cultivars is the same for each trait so that if we order the phenotypic observations of the first trait, then the second trait and so forth, 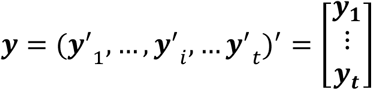 ; then the genetic random effects can be modeled as a normal distribution ***u****∼N*(**0, *U***_*T*_**⊗*K***), where ***U***_*T*_ is a matrix of genomic covariances between the traits of size *t* × *t* to be estimated, and **⊗** indicates the Kronecker product. The matrix ***K*** represents the relationships between the genotypes built with the molecular markers. The random errors are modeled as ***ε****∼N*(**0, Σ⊗*I***), where matrix **Σ** is a matrix of size *t* × *t*, expressing the covariances of the errors between environments to be estimated; and ***I*** is the identity matrix of order *n*_*L*_ × *n*_*L*_. In this study it is assumed that **Σ** is a diagonal matrix that needs to be estimated.

This model MT-SE can also be represented as a multi-response model, that is, instead of outlying the observations as a vector, they can be arranged in a matrix so that *M2* can be re-written as:

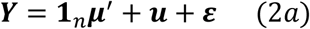

where ***Y*** is a matrix of order *n*_*l*_ × *t* that represents the phenotypic values ordered in such a way that the columns contain the data for each trait and the rows contain the data for each line or genotype. The intercepts or means of each trait are represented by a vector ***µ*** of size *t* × 1. The matrix of genetic random effects assumes that they follow a multivariate multi-response normal distribution 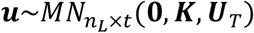. The random errors assume a multivariate multi-response normal distribution 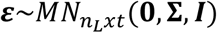, where **Σ** is a matrix of size *t* × *t* denoting the variances-covariances of the random errors within and between traits. In this study we assumed that **Σ** was a diagonal matrix that needs to be estimated.

As already mentioned, when multi-trait data are available, the models to be used are those that account for correlations between phenotypic traits because when the degree of correlation is moderate or large, this could increase the GP accuracy. The model, based on the Bayesian multi-trait kernel of Montesinos et al. (2022), can be seen as the combination of the multi-trait (MT) model *2a* and the reaction norm G×E *M1* for multi-environment (ME). Then *M4* is represented as:

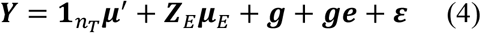

where the matrix ***Y*** is of size *n*_*T*_ × *t* ordered in such a way that the columns represent the phenotypic values of each of the *t* traits and the rows are the lines or genotypes, ordered first by environments and then by lines. The vector ***µ*** is of size *t* × 1 and it represents the intercept or mean of each trait. The matrix ***Z***_*E*_ is an incidence matrix of the environments of size *n*_*T*_ × *m*, and ***µ***_*E*_ is a matrix of order *m* × *t* with the means of each environment in each trait. The matrix ***g*** is of order *n*_*T*_ × *t* and follows a normal distribution 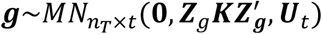 where ***Z***_*g*_ is an incidence matrix of the genotypes of order *n*_*T*_ × *n*_*l*_, ***K*** is the relationship matrix of the genotypes of size *n*_*l*_ × *n*_*l*_ and ***U***_*t*_ is a variance-covariance matrix of traits and between the traits. Matrix ***ge*** is of order *n*_*T*_ × *t* and follows a normal distribution 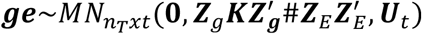 where # is the Hadamard product. Random errors are represented by the matrix ***ε*** of order *n*_*T*_ × *t* that follows a normal distribution 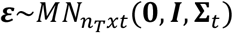 where the identity matrix ***I*** is of dimension *n*_*T*_ × *n*_*T*_ (for more details, see Montesinos et al., 2021).

#### 2.5.5 Study different models and cross-validation schemes to assess the accuracy of the GP prediction models

The GP accuracy of the different models can be assessed by means of several different random cross-validation schemes. The first validation scheme (predicts 100% of the cultivars next year) uses the traits from each of the three locations in 2020 (HEL, MOS, and UM) to predict all the values of the traits in each three locations in 2021 (HEL, MOS, and UM). The second validation scheme (predicts 70% next year) uses all the data from 2020 plus 30% of the value of the traits in three locations in 2021 to predict 70% (prediction set) of the value of the traits at the three locations in 2022; this second case was established with 10 groups or random samples.

A graphical explanation of the different combinations of models (*M1*–M4), considering two prediction sets (100% and 70%), and ST or MT cross-validation schemes for assessing GP prediction accuracy of the models is shown in **Figure 1** for 10 hypothetical cultivars evaluated in HEL, MOS, and UM in 2020 to predict HEL in 2021. The training set (TS) is blue in color and the prediction set (PS) is green. The red lines separate 5 different cross-validation schemes, whereas black lines denote ST prediction, and no lines denote MT predictions. The only MT model is *M4*, whereas ST are models *M1, M2*, and *M3*.

**Figure 1.**
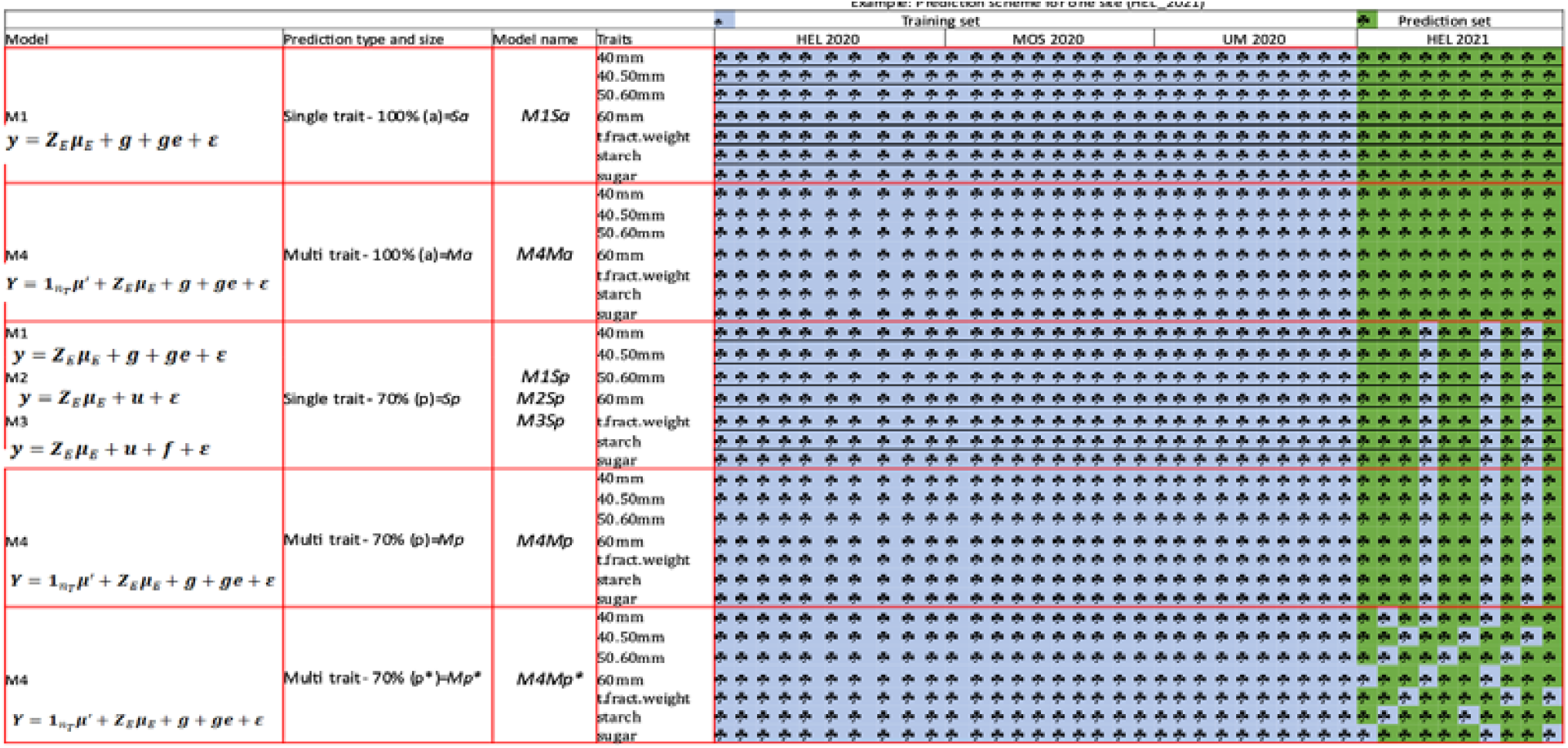
Hypothetical example with 10 cultivars for various models and sizes of prediction set (PS) for the genomic prediction of seven potato traits at Helgegården (HEL) in 2021 (PS) from training data observed at HEL, Mosslunda (MOS) and Umeå (UM) in 2020. Models are *M1*–*M4* and PS are 100% or 70%. The four genome-based prediction models are *M1*: single trait conventional reaction norm model incorporating genomic × environment interaction (GE); *M2*: single trait GE model considering covariances between environments; *M3*: single trait GE M2 extended to include a random vector that more efficiently utilizes the environmental covariances, and *M4*: multi-trait model with GE CV2 is the random cross-validation where 70% are predictive at HEL 2021. Red lines delineated the five random partitions combinations and black lines identified single trait genomic prediction (STGP) and absence of black lines identified multi-trait genomic prediction (MTGP).

As shown in **Figure 1**, the first cross-validations refer to two cases including models M1 and M4 for predicting all the values (100%) for each trait in location HEL 2021 using as a training set all the values for each trait in each location from 2020. Model *M1* is an ST (traits are separated by black lines), whereas *M4* is an MT model (traits are not separated). For these two cases, the given names join (1) the model, (2) the ST or MT (S or M) type of prediction, and (3) include the prediction of all (100%) the lines in HEL 2021 and denoted by ‘a’, that is, *M1Sa* and *M4Ma*. The third and fourth cross-validation schemes delineated by red lines included models *M1, M2, M3* for ST and model *M4* for MT, and they predict 70% of the values of each trait in HEL 2021 using as training set values of the trait in each location from 2020 but also adding 30% of the values from HEL 2021 to the prediction set in the training set. As already mentioned, this prediction of 70% is performed 10 times using the 10 random samples for extracting 30% of the values of the prediction set (2021) and adding them into training set (2020). The same 10 random samples were used for comparing the genomic prediction accuracy of the four models. The names of each of these model-prediction types and sizes are *M1Sp, M2Sp, M3Sp*, and *M4Mp* where the letter ‘*p’* refers to the percentage of the prediction set (70%). Note that for these four cases, 3 cultivars (out of 10) are missing in all the traits (**Figure 1**). The fifth cross-validation scheme had MT *M4* that predicts 70% of the cultivars in HEL in 2021 for all traits but now the cross-validations between the traits and locations for HEL 2021 are different from those in the previous case (*M4Mp*) where some cultivars are observed in some traits and locations but not observed in other traits and locations. This cross-validation scheme is refereed to *M4Mp\** Note that in this case, some cultivars are missing in some traits but not in other traits; for example, cultivars 1, 2, and 3 are not observed for weight of tubers below 40 mm but are observed for the weight of 40−50 mm tubers (**Figure 1**).

#### 2.5.6 Measures of prediction accuracy

We used two metrics for comparing the genomic-enabled prediction accuracy of the different models (*M1, M2, M3*, and *M4*). One metric is the Pearson correlation coefficient (COR) between the observed and predicted values, whereas the second metric is the prediction mean squared error (PMSE) of the different prediction models.

## 3. RESULTS

Phenotypic correlations were computed for traits in each location (HEL, MOS, and UM) in 2021 (PS) with those traits in the locations of the previous year (HEL, MOS, and UM in 2020) (**Table 1**). The PS contains seven traits (5 tuber weight traits and 2 tuber flesh quality characteristics) in each of the 3 locations of 2021 using the locations and traits of the previous year, 2020. The ST or MT prediction models together with the proportion of cultivars included in the PS are combined in *M1Sa, M4Ma, M1Sp, M2Sp, M3Sp, M4Mp*, and *M4Mp\** (**Tables 2−4** and **Figures 2−4**).

**Table 1.** Phenotypic correlations of each trait at Helgegården (HEL) in 2021 with each trait at HEL 2020, Mosslunda (MOS) 2020, and Umeå (UM) 2020. Phenotypic correlations of each trait oat MOS 2021 with each trait at HEL 2020, MOS 2020, and UM2020. Phenotypic correlations of each trait at UM 2021 with each trait at HEL 2020, MOS 2020, UM 2020.

| Site_year | Traits |  |  |  |  |  |  |
| --- | --- | --- | --- | --- | --- | --- | --- |
|  | Weight of tubers |  |  |  |  | Tuber flesh |  |
|  | <40 mm | 40–50 mm | 50–60 mm | > 60mm | Total | Starch | Sugar |
| HEL 2021 |  |  |  |  |  |  |  |
| HEL 2020 | 0.62 | 0.60 | 0.24 | 0.68 | 0.64 | 0.89 | 0.36 |
| MOS 2020 | 0.36 | 0.20 | -0.16 | 0.49 | 0.48 | 0.80 | 0.30 |
| UM 2020 | 0.43 | -0.05 | -0.25 | 0.51 | 0.39 | 0.78 | 0.43 |
| MOS 2021 |  |  |  |  |  |  |  |
| HEL 2020 | 0.65 | 0.49 | 0.56 | 0.73 | 0.64 | 0.83 | 0.39 |
| MOS 2020 | 0.64 | 0.50 | 0.61 | 0.74 | 0.74 | 0.89 | 0.36 |
| UM 2020 | 0.55 | 0.28 | 0.45 | 0.62 | 0.52 | 0.72 | 0.41 |
| UM 2021 |  |  |  |  |  |  |  |
| HEL 2020 | 0.49 | 0.04 | 0.42 | 0.53 | 0.38 | 0.48 | 0.31 |
| MOS 2020 | 0.49 | 0.30 | 0.47 | 0.40 | 0.29 | 0.40 | 0.33 |
| UM 2020 | 0.57 | 0.51 | 0.67 | 0.57 | 0.46 | 0.46 | 0.46 |

**Table 2.**
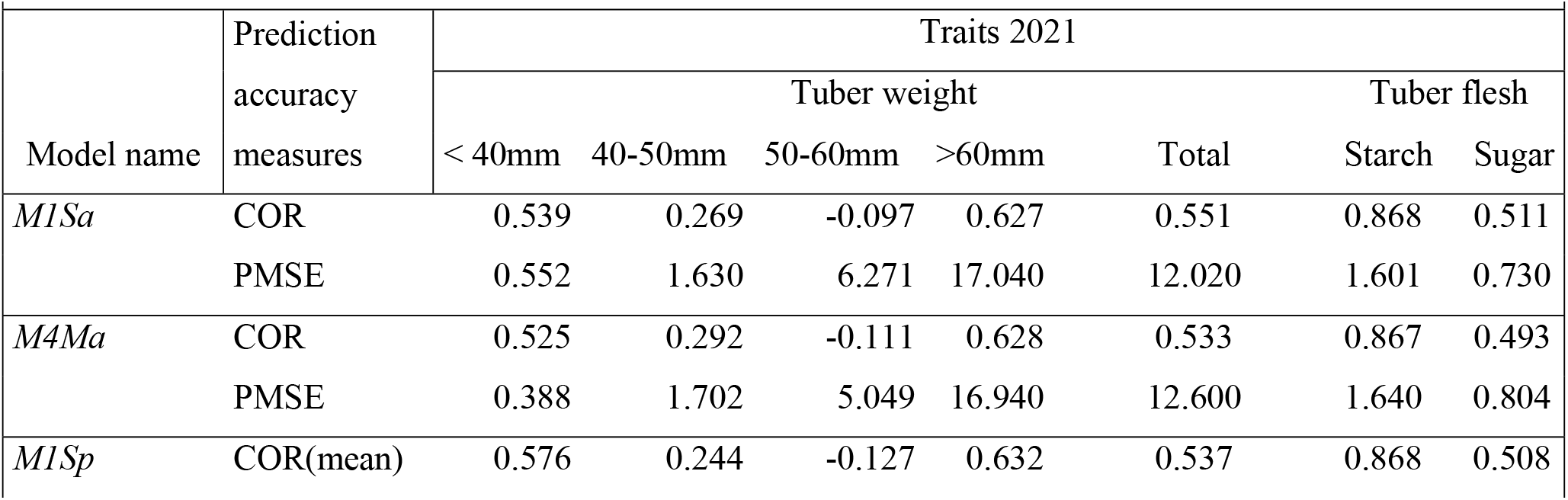
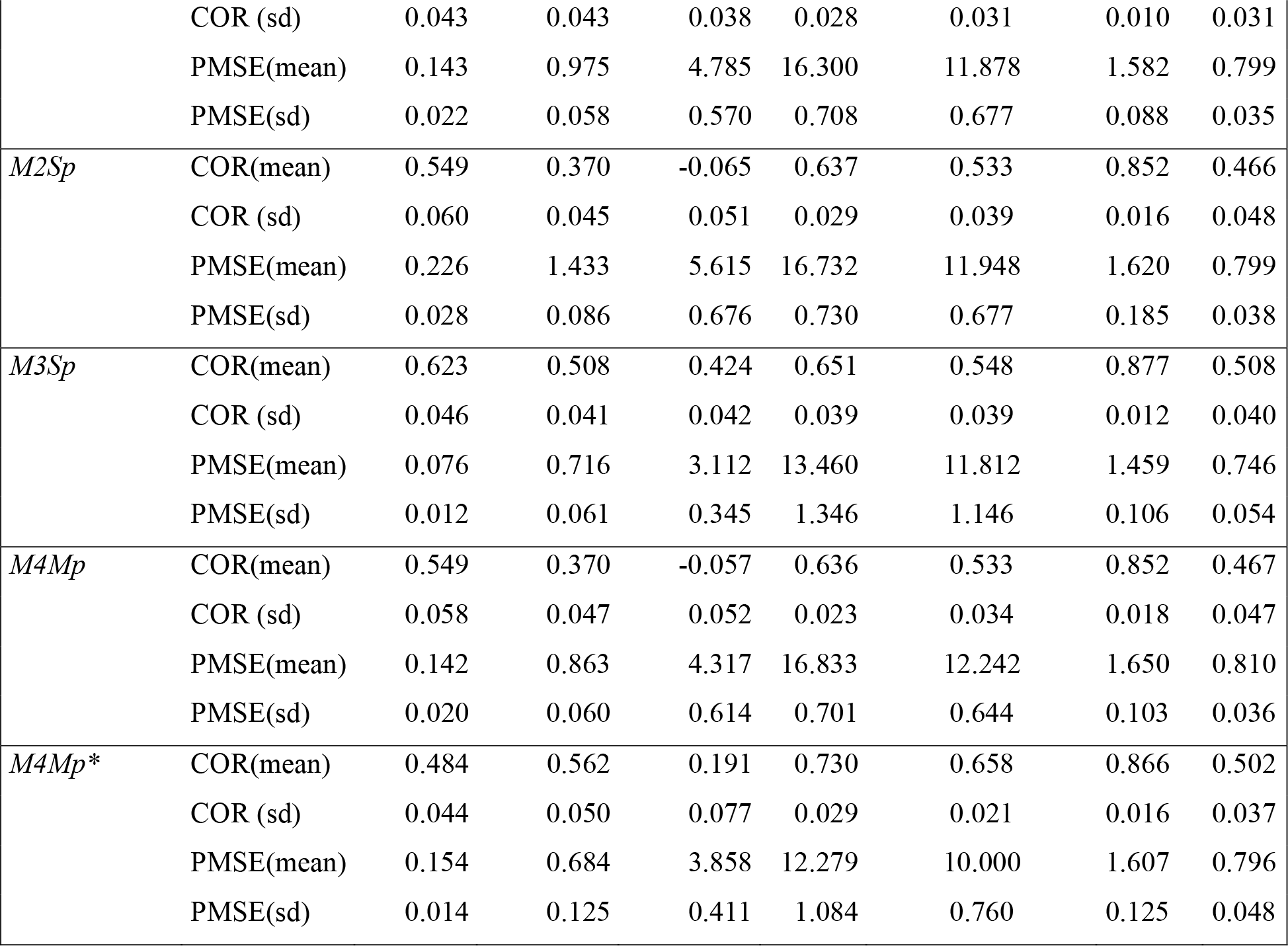
Predictive correlations (COR) and predictive mean squared error (PMSE) for predicting seven traits at Helgegården (HEL) in 2021 for four models (*M1, M2, M3, M4*) combined with 100% or 70% cross-validation. *M1Sa* is the prediction accuracy from model *M1* (single trait conventional reaction norm model incorporating genomic × environment interaction [GE]) when predicting 100% of each trait in 2021; *M4Ma* is the prediction accuracy from model *M4* (multi-trait model with GE) when predicting 100% of each trait in 2021; *M1Sp* is the prediction accuracy from model *M1* when predicting 70% of each trait in 2021; *M2Sp* is the prediction accuracy from model *M2* (single trait GE model considering covariances between environments) when predicting 70% of each trait in 2021; *M3Sp* is the prediction accuracy from model *M3* (single trait GE *M2* extended to include a random vector that more efficiently utilizes the environmental covariances) when predicting 70% of each trait in 2021; *M4Mp* is the prediction accuracy from model *M4* when predicting 70% of each trait in 2021, *M4Mp\** is the prediction accuracy from model *M4* when predicting 70% of each trait in 2021 in which some cultivars are observed in some traits. When predicting 70%, the mean and the standard deviations (sd) from the 10-fold cross-validation are given in parentheses.

**Figure 2.**
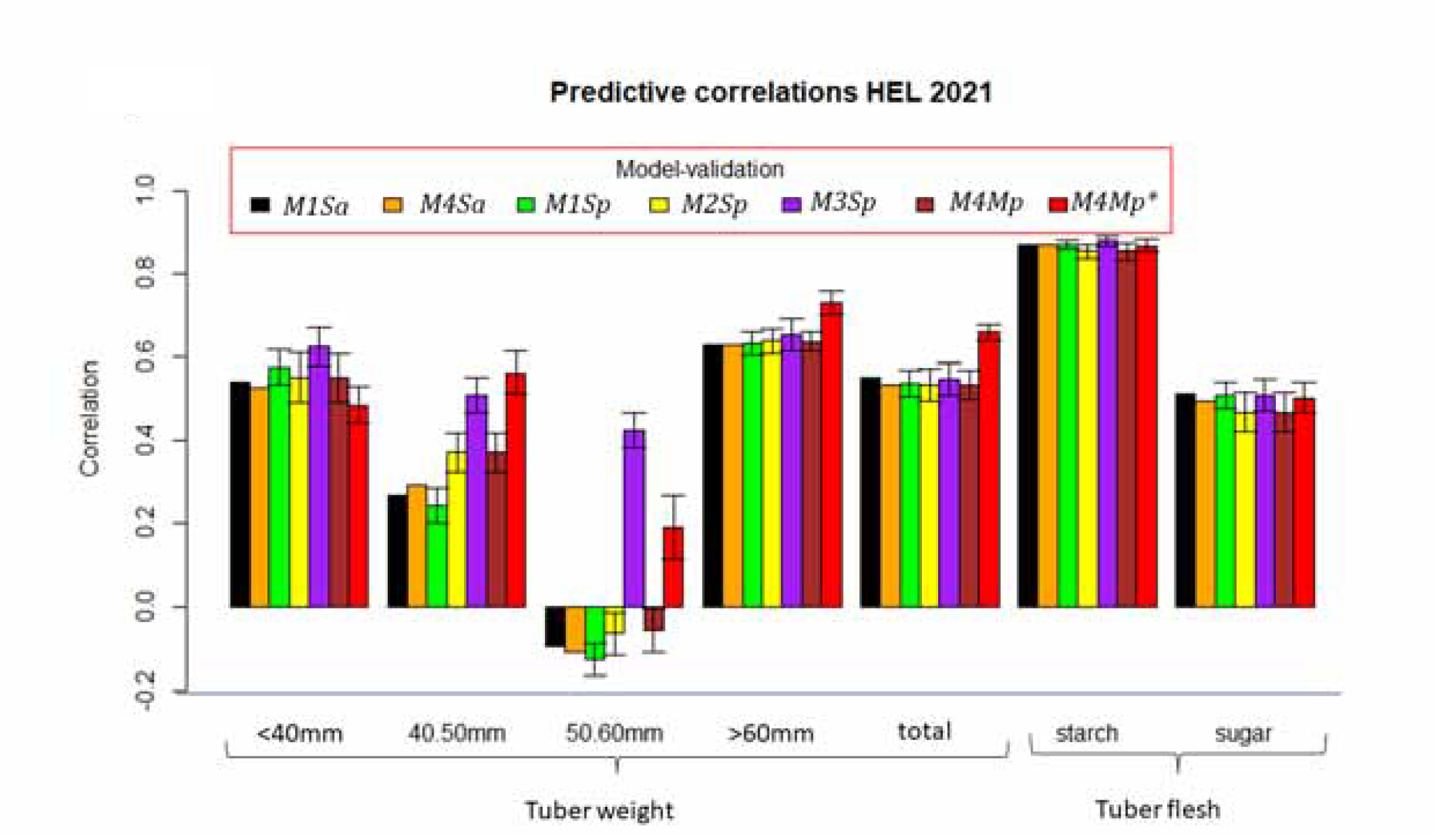
Trait prediction in 2021 at Helgegården (HEL). *M1Sa* is the prediction accuracy from model *M1* (single trait conventional reaction norm model incorporating genomic × environment interaction [GE]) when predicting 100% of each trait in 2021). *M4Ma* is the prediction accuracy from model *M4* (multi-trait model with GE) when predicting 100% of each trait in 2021. *M1Sp* is the prediction accuracy from model *M1* when predicting 70% of each trait in 2021. *M2Sp* is the prediction accuracy from model *M1* when predicting 70% of each trait in 2021. *M3Sp* is the prediction accuracy from model *M1* when predicting 70% of each trait in 2021. *M4Mp* is the prediction accuracy from model *M4* when predicting 70% of each trait in 2021. *M4Mp\** is the prediction accuracy from model *M4* when predicting 70% of each trait in 2021 in which some cultivars are observed in some traits.

### 3.1 Genomic prediction of traits in HEL 2021

Results are presented by location−year combination and predictions included the whole location in year 2021 (*M1Sa*, and*M4Ma*) and prediction of only 70% of the 2021 location (*M1Sp, M2Sp, M3Sp, M4Sp*, and *M4Sp\**). Phenotypic correlations of traits measured in HEL, MOS, and UM 2020 with all the traits measured in HEL-2021 are given in **Table 1**. The phenotypic correlations between traits in HEL for 2020 and 2021 are higher than those between HEL 2021 and other locations in 2020. Tuber flesh starch had the highest phenotypic correlation between HEL 2021 and HEL, MOS, and UM 2020 (0.89, 0.80, and 0.78, respectively) followed by weight of tubers above 60 mm (0.68, 0.49, and 0.51, respectively), total tuber weight irrespective of size (0.64, 0.48, and 0.39, respectively), and weight of tubers below 40 mm (0.62, 0.36, and 0.43, respectively).

Genomic predictions of whole traits in HEL 2021 from *M1Sa* and*M4Ma* as well as from *M1Sp* to*M4Mp\**, clearly show tuber flesh starch as the best predicted trait for all the models with genomic prediction accuracy above 0.85 (**Table 2** and **Figure 2**). Most of the four models shown a very similar genomic prediction accuracy for trait starch ranging from 0.852 (*M2Sp* and *M4Mp*) to 0.877 (*M3Sp*) (**Table 2, Figure 2**).

The second trait with important GP accuracy shown by most of the models was tuber weight of 60 mm where *M4Mp\** had the highest prediction accuracy (0.730, **Table 2**) and *M1Sa* had the lowest genomic prediction accuracy (0.627). Weight of tubers below 40 mm and total tuber weight had very similar genomic prediction accuracy except for model *M4Mp\** which was the worst model for weight of tubers below 40 mm but the best model for trait total tuber weight. Excluding *M4Mp\**, the predictions ranged from 0.525 (<40 mm, *M4Ma*) to 0.623 (<40mm *M3Sp*) for both traits. The best predictive model was *M3Sp* for weight of tubers below 40 mm and *M1Sa* for total tuber weight (**Figure 2**). Weight of tubers with 40–50 mm and 50–60 mm sizes had the lowest prediction accuracy for most models except *M3Sp* (**Figure 2**). Comparing models with ST and MT, *M3Sp* was the best ST model for 3 traits (tuber weight below 40mm and between 50–60mm, and tuber flesh starch) and *M4Mp\** was best for the other 3 traits (40–50mm, >60mm, and total tuber weight).

In summary, prediction of the seven traits at HEL in 2021 shows that traits with higher phenotypic correlation between location HEL 2021 and those at HEL, MOS, and UM in 2020 are tuber flesh starch and most of the tuber weights (except weight of tubers 50–60 mm). In terms of GP accuracy, multi-trait model *M4Mp\** was the best in weight of tubers 40−50mm or above 60 mm size, and total tuber weight, being very similar to those for tuber flesh starch. Model *M3Sp* was the best GP for tuber weights <40mm and 50– 60mm, as well as tuber flesh starch.

### 3.2 Genomic prediction of traits in MOS 2021

Phenotypic correlation of traits measured in location MOS in 2020–2021 are given in **Table 1**. For all the traits the phenotypic correlations between traits in MOS for 2021 and 2020 are higher than those between MOS 2021 and the two other locations (HEL and UM) in 2020. Tuber flesh starch had the highest phenotypic correlation between MOS 2021and HEL, MOS, and UM 2020 (0.83, 0.89, and 0.72, respectively) followed by weight of tubers above 60 mm (0.73, 0.74, and 0.62, respectively), total tuber weight (0.64, 0.74, and 0.52, respectively), and weight of tubers below 40 mm (0.65, 0.64, and 0.55, respectively).

Overall genomic predictions accuracy in MOS 2021 was higher than in HEL 2021. Tuber flesh starch was the best predicted trait for all the models with < 0.85 genomic prediction accuracy (**Table 3** and **Figure 3**). Most of the four models showed a very similar genomic prediction accuracy for tuber flesh starch but *M2Sp* and *M3Sp* were the best genomic predictors, with 0.866 and 0.867, respectively. *M1Sa* and*M4Ma* were slightly below in terms of prediction accuracy (0.847 and 0.848, respectively).

**Table 3.**
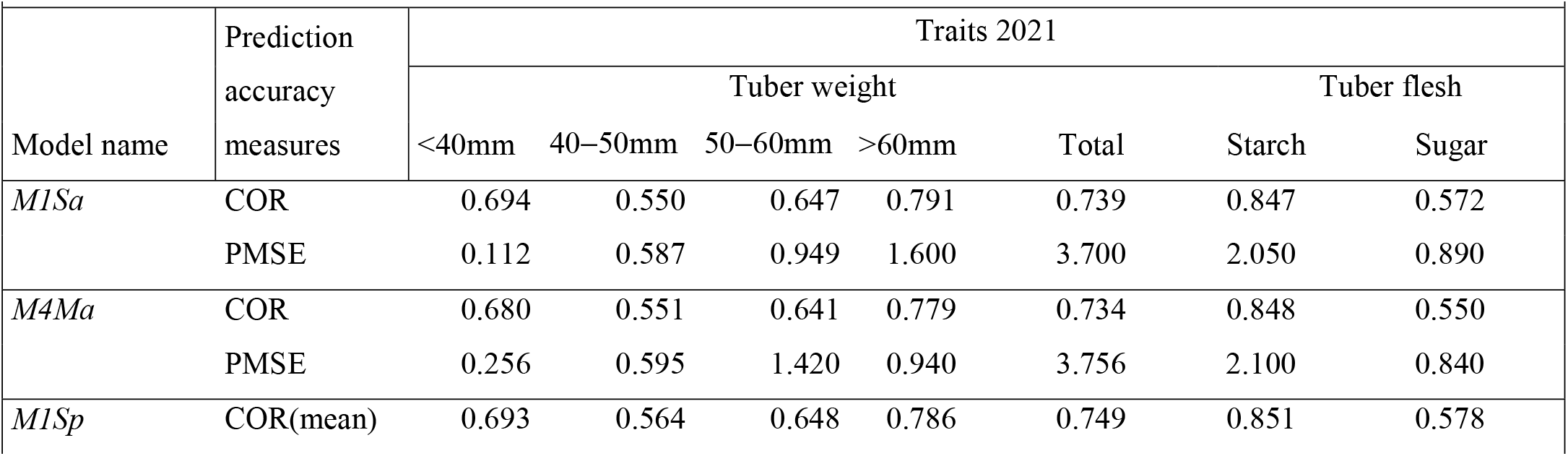
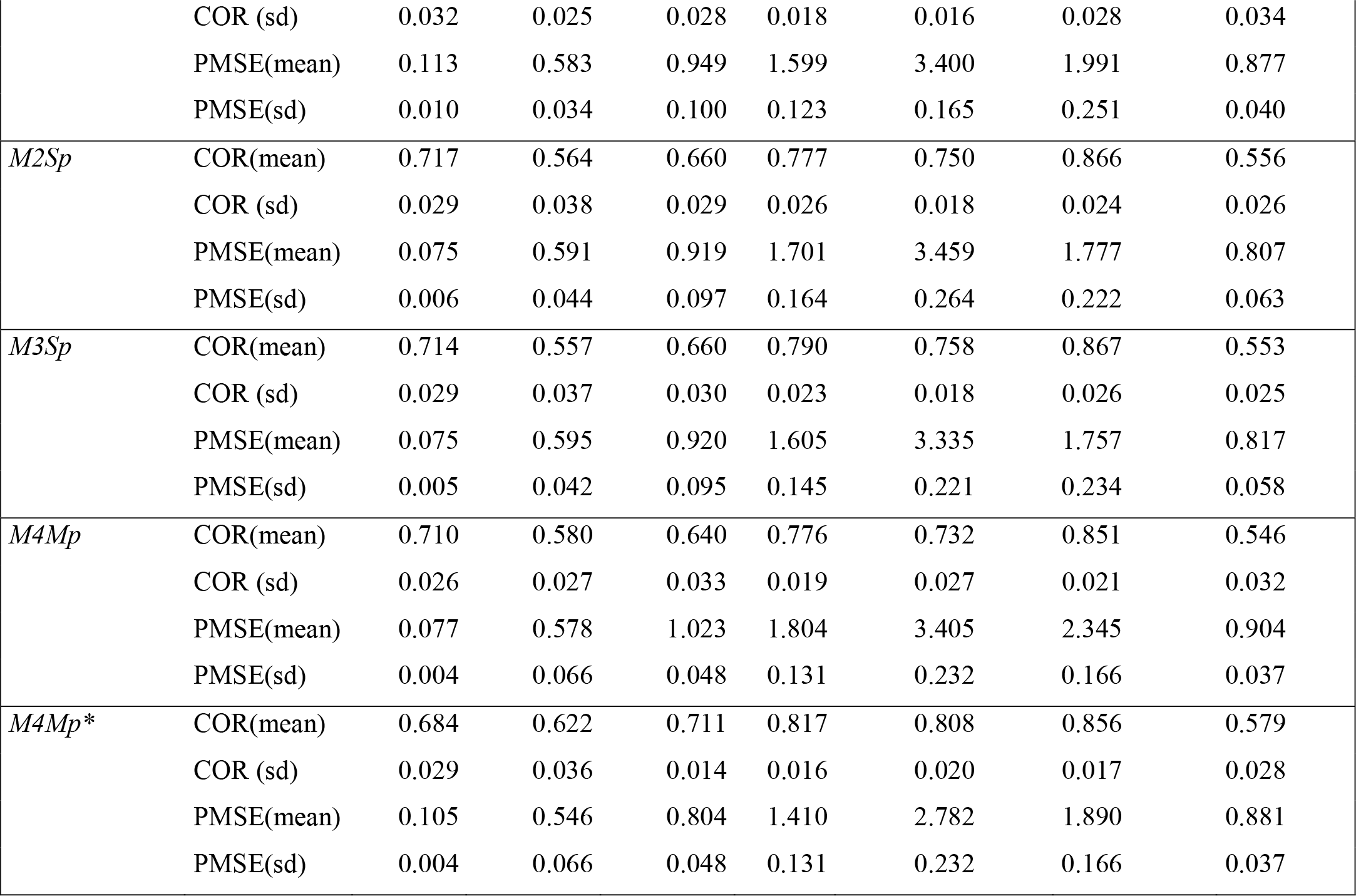
Predictive correlations (COR) and predictive mean squared error (PMSE) for predicting seven traits at Mosslunda (MOS) in 2021 for four models (*M1, M2, M3, M4*) combined with 100% or 70% cross-validation. *M1Sa* is the prediction accuracy from model *M1* (single trait conventional reaction norm model incorporating genomic × environment interaction [GE]) when predicting 100% of each trait in 2021. *M4Ma* is the prediction accuracy from model *M4* (multi-trait model with GE like) when predicting 100% of each trait in 2021. *M1Sp* is the prediction accuracy from model *M1* when predicting 70% of each trait in 2021. *M2Sp* is the prediction accuracy from model *M2* (single trait GE model considering covariances between environments) when predicting 70% of each trait in 2021. *M3Sp* is the prediction accuracy from model *M3* (single trait GE *M2* extended to include a random vector that more efficiently utilizes the environmental covariances) when predicting 70% of each trait in 2021; *M4Mp* is the prediction accuracy from model *M4* when predicting 70% of each trait in 2021, *M4Mp\** is the prediction accuracy from model *M4* when predicting 70% of each trait in 2021 in which some cultivars are observed in some traits. When predicting 70%, the mean and the standard deviations (sd) are given from the 10-fold cross-validation in parentheses.

**Figure 3.**
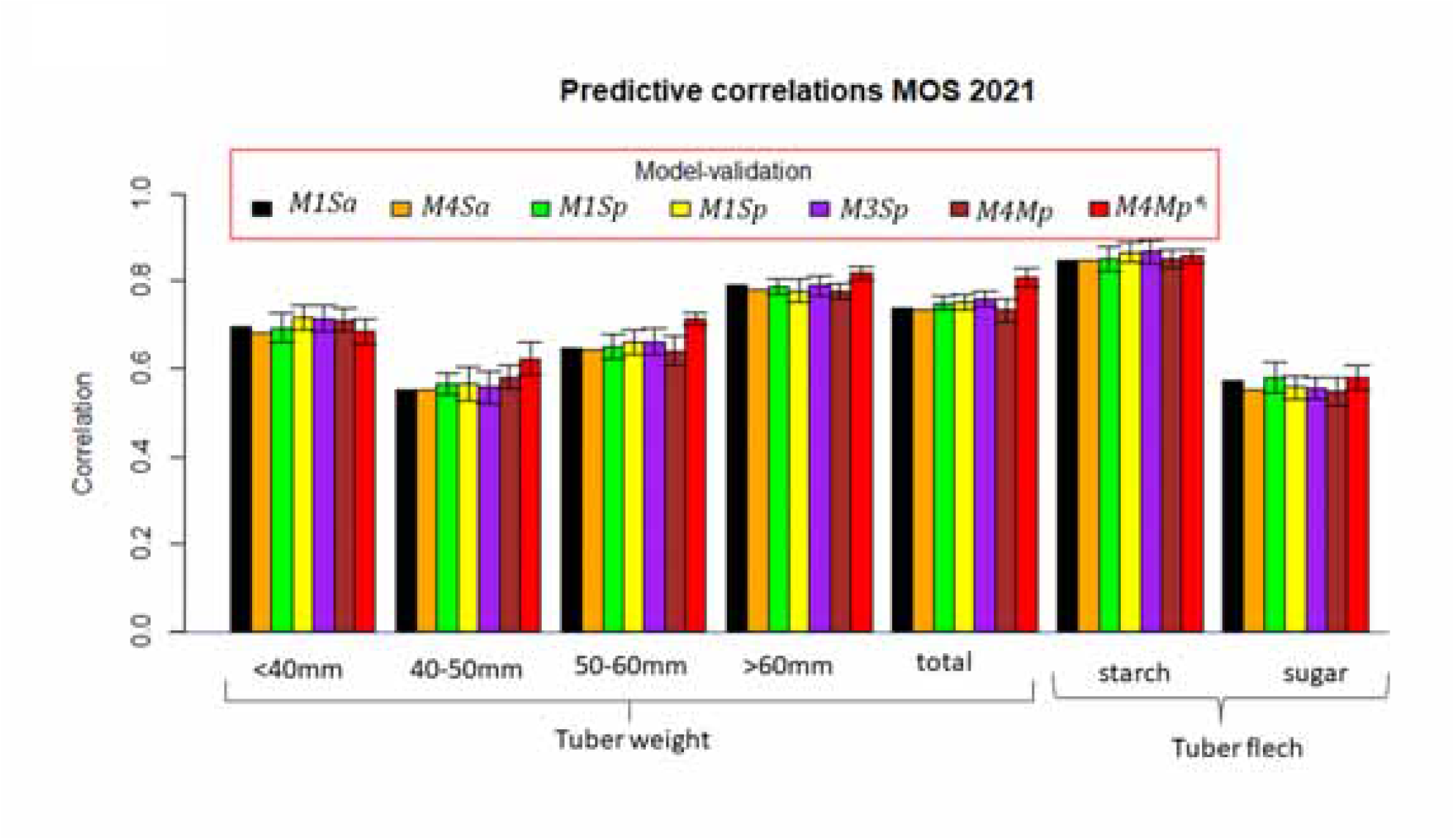
Trait prediction in 2021 at Mosslunda (MOS). *M1Sa* is the prediction accuracy from model *M1* (single trait conventional reaction norm model incorporating genomic × environment interaction [GE]) when predicting 100% of each trait in 2021. *M4Ma* is the prediction accuracy from model *M4* (multi-trait model with GE) when predicting 100% of each trait in 2021. *M1Sp* is the prediction accuracy from model *M1* when predicting 70% of each trait in 2021. *M2Sp* is the prediction accuracy from model *M1* when predicting 70% of each trait in 2021. *M3Sp* is the prediction accuracy from model *M1* when predicting 70% of each trait in 2021. *M4Mp* is the prediction accuracy from model *M4* when predicting 70% of each trait in 2021. *M4Mp\**is the prediction accuracy from model *M4* when predicting 70% of each trait in 2021 in which some cultivars are observed in some traits.

The second trait with important genomic prediction accuracy shown by most of the models was tuber weight above 60 mm with *M4Mp\** with an accuracy of 0.817, followed by*M1Sa* having an accuracy of 0.791 followed by *M3Sp* with 0.790 (**Table 3**). Overall, total tuber weight irrespective of size ranked third based on genomic prediction accuracy, with model *M4Mp\** having a prediction accuracy of 0.808, followed by *M3Sp* with 0.758 prediction accuracy followed by *M2Sp* (0.750). Weight of tubers below 40mm had relatively high genomic prediction accuracy, with models *M2Sp* and *M3Sp* being the best with 0.717 and 0.714 of genomic prediction accuracy, respectively. Finally, weight of tubers 50−60 mm in size had lower prediction accuracy than the previously mentioned traits, with the best predictor models being *M4Mp\** with 0.711 GP accuracy, followed by *M2Sp* and *M3Sp* with 0.660 accuracy.

The GP accuracy of the seven traits in location MOS in 2021 showed slightly higher accuracy in the prediction of the seven traits in 2021 than those found for the traits at HEL 2021. Results show that the traits with higher phenotypic correlation between MOS 2021 and those at HEL, MOS, and UM in 2020 are tuber flesh starch, weight of tubers above 60 mm and below 40 mm, total tuber weight, and weight of tubers with 50–60mm. In general, the best models for predicting the majority of the seven traits were *M3Sp* and *M2Sp*, except for traits such as weight of tubers with 50–60mm and above 60 mm, and total tuber weight in which MT model *M4Mp\** was the best GP model.

### 3.3 Genomic prediction of traits in location UM 2021

**Table 1** lists the phenotypic correlation of traits measured at UM in 2020−2021. For all the traits, the phenotypic correlations between traits in UM for 2021 and 2020 are higher than those between UM 2021 and other locations (HEL and MOS) in 2020. The traits with the highest phenotypic correlation between UM 2021 and HEL, MOS and UM 2020 were weight of tubers with 50−60mm, below 40 mm, and above 60 mm, followed by tuber flesh starch.

Overall genomic prediction accuracy in UM 2021 was lower than those found at HEL and MOS in 2021. Weight of tuber with 50−60 mm and below 40 mm were the best predicted traits for all the models in UM 2021 (**Table 4** and **Figure 4**). The best GP model for all the traits, except reducing sugars and starch in the tuber flesh, was *M4Mp\**. Models *M3Sp* and *M4Mp* had the best GP accuracy for predicting traits tuber flesh sugar and starch, respectively.

**Table 4.**
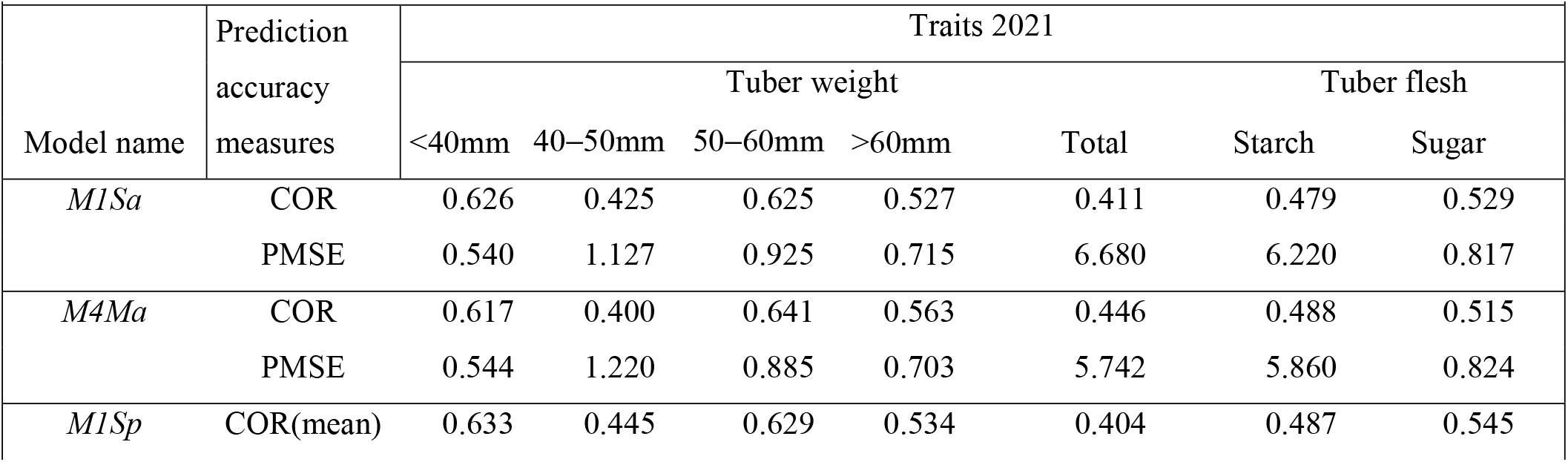
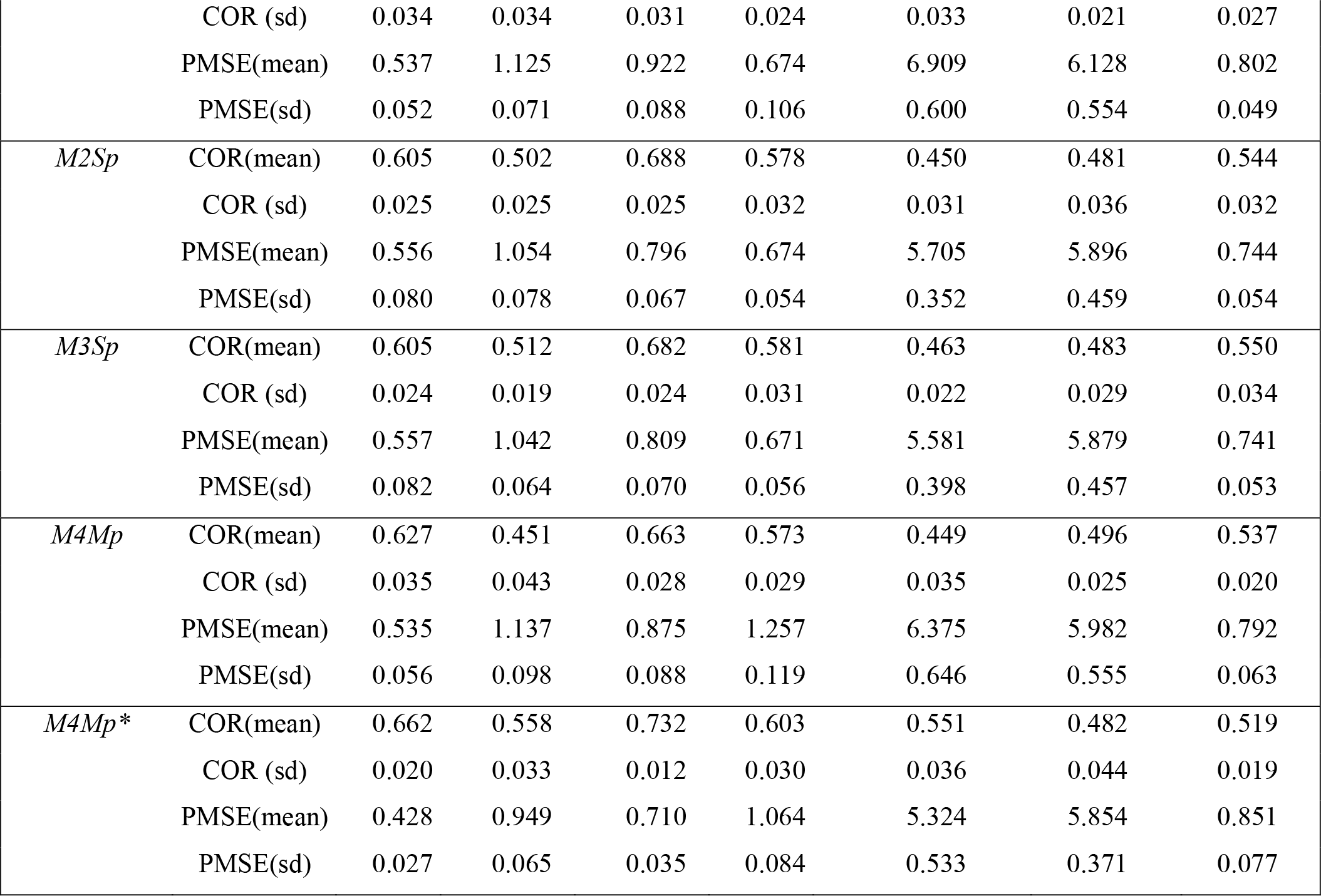
Predictive correlations (COR) and predictive mean squared error (PMSE) for predicting seven traits at UM in 2021 for four models (*M1, M2, M3, M4*) combined with 100% or 70% cross-validation. *M1Sa* is the prediction accuracy from model *M1* (single trait conventional reaction norm model incorporating genomic × environment interaction [GE]) when predicting 100% of each trait in 2021. *M4Ma* is the prediction accuracy from model *M4* (multi-trait model with GE) when predicting 100% of each trait in 2021. *M1Sp* is the prediction accuracy from model *M1* when predicting 70% of each trait in 2021. *M2Sp* is the prediction accuracy from model *M2* (single trait GE model considering covariances between environments) when predicting 70% of each trait in 2021. *M3Sp* is the prediction accuracy from model *M3* (single trait GE *M2* extended to include a random vector that more efficiently utilizes the environmental covariances) when predicting 70% of each trait in 2021. *M4Mp* is the prediction accuracy from model *M4* when predicting 70% of each trait in 2021, *M4Mp\** is the prediction accuracy from model *M4* when predicting 70% of each when some cultivars are observed in some traits. When predicting 70% the mean and the standard deviations (sd) from the 10-fold cross-validation are given in parentheses.

**Figure 4.**
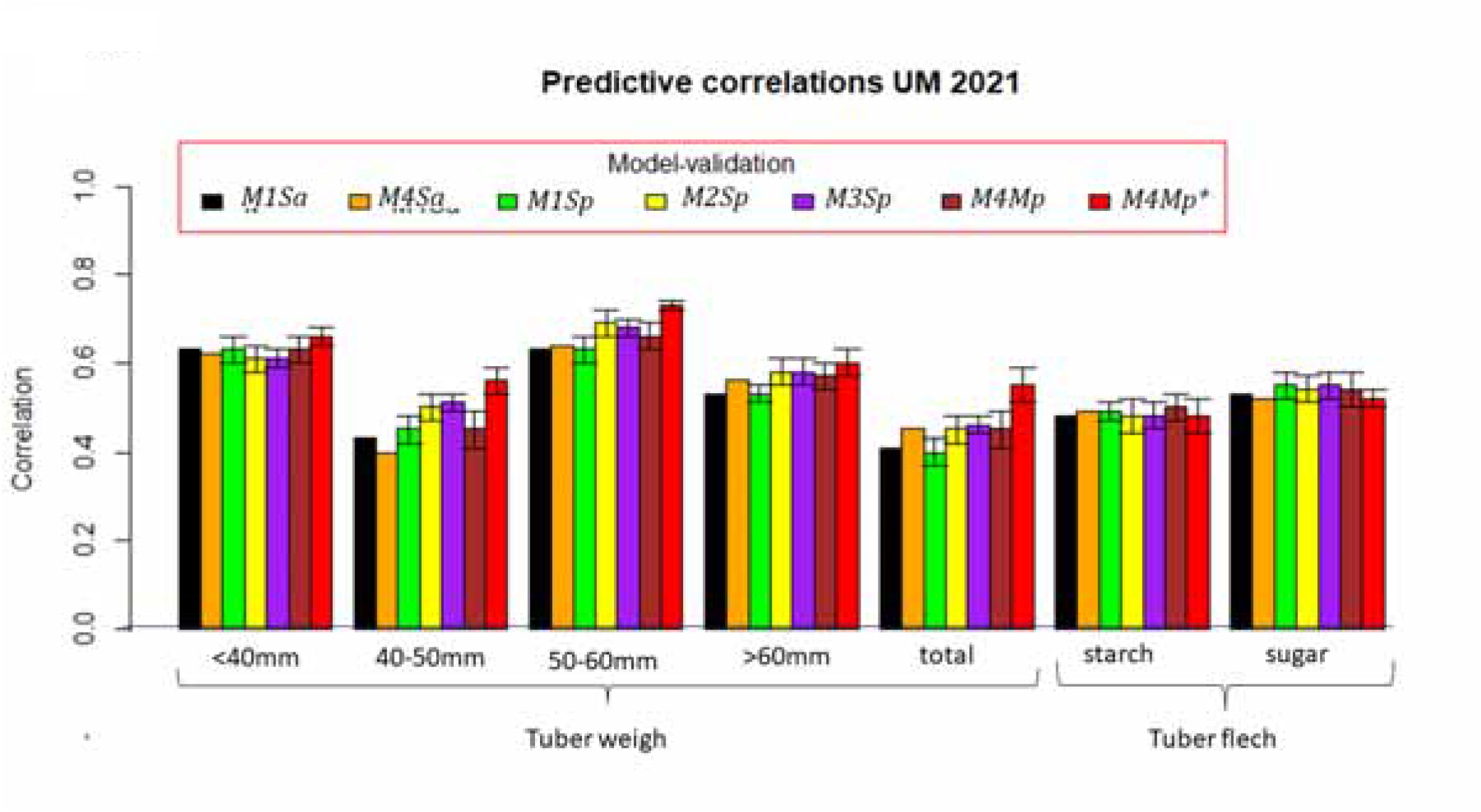
Trait prediction in 2021 at Umeå (UM). *M1Sa* is the prediction accuracy from model *M1* (single trait conventional reaction norm model incorporating genomic × environment interaction [GE]) when predicting 100% of each trait in 2021. *M4Ma* is the prediction accuracy from model *M4* (multi-trait model with GE) when predicting 100% of each trait in 2021. *M1Sp* is the prediction accuracy from model *M1* when predicting 70% of each trait in 2021. *M2Sp* is the prediction accuracy from model *M1* when predicting 70% of each trait in 2021. *M3Sp* is the prediction accuracy from model *M1* when predicting 70% of each trait in 2021. *M4Mp* is the prediction accuracy from model *M4* when predicting 70% of each trait in 2021. *M4Mp\** is the prediction accuracy from model *M4* when predicting 70% of each trait in 2021 in which some cultivars are observed in some traits

Most of the four models showed similar genomic prediction accuracy for these two traits, but *M2Sp* had a genomic prediction accuracy of 0.688 for tuber weight with 50−60mm and model *M4Mp* had an accuracy of 0.633 for weight of tubers below 40 mm. Models *M2Sp* and *M3Sp* had a genomic prediction accuracy of around 0.578 for weight of tubers above 60 mm that ranked third on overall genomic prediction accuracy (**Table 4**) followed by tuber flesh starch, with model *M3Sp* being the best with 0.483 prediction accuracy, followed by *M2Sp* (0.481).

The genomic prediction accuracy of the seven traits at UM in 2021 showed lower accuracy in 2021 than at HEL and MOS in 2021. Traits with higher phenotypic correlations between UM 2021 and those at HEL, MOS, and UM in 2020 are weight of tubers with 50−60mm, below 40 mm, and above 60 mm. However, the best model for predicting the majority of the seven traits was *M4Mp\**, followed by models *M4Mp* for tuber flesh starch and *M3Sp* for tuber flesh sugar.

## 4. DISCUSSION

The integration of GS and GP to develop modern cultivars faster than the conventional breeding method is necessary for increasing genetic gains and facing the changes in climate that are currently affecting agriculture. Thus, a better and efficient integration of new methods including GS with increased GP accuracy, rapid cycle GS, high throughput phenotyping, and the use of appropriate environmental covariables is an urgent area of research (Crossa et al., 2021). The integration and exploitation of several big data sets is necessary, and the use of appropriate statistical machine learning models has become important for modern breeding.

### 4.1 Prediction accuracy of model for ST and MT, cross-validation method and proportion of the prediction set

When performing research on GS and GP accuracy, several problems become important; one is the inclusion of statistical machine learning methods and models that include GE interaction. Another problem to be assessed is the addition of several traits for prediction rather than only one trait, and another issue is the methods used for comparing the GP accuracy of several traits using several models and various possible cross-validation schemes to develop a GP accuracy metric. Several options exist for investigating the GS accuracy for predicting the breeding value of cultivars that have been genotyped with genome-wide molecular markers. One scenario is predicting the performance of a proportion of cultivars (e.g., 70%) that have not yet been observed in any of the testing environments (usually location-year combinations); another option is to predict all cultivars (i.e., 100%) observed in all the environments except one (leave one environment out). Another scenario is predicting cultivars that were observed in some environments but not in others.

In this study predictions for these scenarios have been done using single-trait (ST) (*M1, M2* and *M3*) and multi-trait (MT) (*M4*) models. These ST and MT models combined with different PT scenarios are represented in **Figure 1**, where several proportions of the PS have been combined with the four different models. We included the predictions of all cultivars in one entire site-year combination or the prediction of a proportion of cultivar (70%) using the other 30% as TS together with the previous year. We found that for the majority of the traits in each location-year combination to be predicted (HEL, MOS, UM in 2021) *M4* (multi-trait), with a proportion of potato cultivars evaluated (30%) in some location-year combinations *M4Mp\** (**Figure 1**) but not observed in other location-year combinations, was found to be the best predictive model usually followed by ST models *M3Sp* and *M2Sp*.

Results of this study demonstrate that for predicting traits in HEL 2021 using all environments in 2020 the superiority of the MT prediction method *M4Mp\** over the mean GP accuracy of the other six prediction methods including ST and MT for predicting entire PS (100%) or 70% for traits tuber weights 40–50mm, above 60mm and total in location were 65%, 14% and 24%, respectively. However, this superiority of the MT over ST methods was not so when comparing *M4Ma* or *M4Mp* with other ST methods, especially for *M3Sp* for traits tuber weight < 40mm, 50–60mm and tuber flesh starch. Results for predicting traits in location MOS in 2021 using all environments in 2020 show the superiority of MT prediction method *M4Mp\** for four tuber weight traits and one tuber flesh quality characteristic over all the other six methods. The GP accuracy of method *M4Mp*\* overcame the mean GP accuracy of all the other six methods by 10%, 9%, 4%, 8% and 4% for traits tuber weights 40–50mm, 50–60mm, above 60mm, total and tuber flesh sugar, respectively. Similar results were obtained for the prediction of location UM in 2021 using the TS comprising HEL, MOS, and UM from 2020; the best GP accuracy method for all five tuber weight traits was method *M4Mp\** over the mean GP accuracy of all the other six methods by 7%, 24%, 12%, 8% and 26% for tuber weights below 40 mm, 40–50 mm, 50–60 mm, above 60 mm and total tuber weight, respectively.

Previous research noticed variable prediction accuracy that depends on factors such as heritability of the trait, size of TP, relatedness of PS and TS, statistical machine learning models, marker density, linkage disequilibrium, and the incorporation of GE interactions in the prediction models. In a recent article, Semagn et al. (2022) compared the predictive abilities of wheat cultivars that have not been evaluated for a single trait (ST), not evaluated for multi-traits (MT1), and evaluated for some traits but not others (MT2) using agronomy and disease traits. Note that the partition of Semagn’s MT1 is similar to the partitions of *Sp* (*M1, M2*, and *M3*) and *Mp* (*M4*) in this study, whereas the partitions of Semagn’s MT2 is similar to that of *M4Mp\**. Semagn et al. (2022) found that the GP accuracy of MT2 (method *M4Mp\** in this study) increased over ST and other model-partitions in all traits from 9% to 82%. This occurred because under the prediction scheme MT2 of Semagn et al. (2022) it is possible exchange of information between traits like method *M4Mp\** that allows borrowing of information between traits and also between environments and thus efficiently use the available information in one single model combined with an appropriate prediction scheme.

This demonstrated the high potential for improving prediction accuracies and the high potential of the MT models for improving prediction accuracy, thus offering researchers the opportunity to predict traits that were not observed, due to possible difficulties or because they are expensive to measure under certain environmental constraints (Semagn et al., 2022).

### 4.2 Prediction accuracy of potato traits

Genomic prediction in potato is still in the early research stages before using it for routine breeding of this highly heterozygous tetrasomic polyploid tuberous crop with vegetative propagation (Ortiz et al., 2022, and references therein). The use of MT and ME models for genomic prediction in this research led to highest accuracy for tuber yield and tuber flesh starch as per available literature. Tuber flesh starch, which is often estimated from specific gravity measurements, is a very highly heritable trait (Bradshaw, 2021, Ortiz et al., 2021) that is affected very little by the genotype × environment interactions (Killick and Simmonds, 1974), thus explaining the high prediction accuracy noted in this and research elsewhere. The high prediction accuracy noted in this, and previous research suggest that developing GEBV modeling in potato for tuber flesh starch does not require a very large training population but just a few hundred (including both breeding clones and released cultivars that are relevant to the breeding program and covering a broad range of trait variation) may suffice.

Genotype × environment interactions may significantly affect tuber yield, but the use of multi-environment genomic prediction allows identifying promising germplasm in both crossing blocks (Ortiz et al., 2022) in potato breeding. The significantly high correlations noted when using multi-trait, multi-environment modeling suggest that genomic prediction may also be useful for the potato cultivar development pipeline even when using small breeding populations (Selga et al., 2022). Every year F_1_ seeds (resulting from crossing heterozygous parents) are planted in individual pots in a greenhouse, and one tuber (the best in size) for each plant is taken at harvest. Thus, thousands of tubers derived from these F_1_ hybrid seeds are produced for further field testing in single plant plots during the first year. At harvest, all plants are dug up to assess their tuber number, size, shape, color, appearance, and health, which are used as the selection criteria for obtaining the next breeding generation for further testing the next year. After selection in early clonal generations (first [T_1_], second [T_2_] and often third [T_3_]), the aim is to have about a few dozens for field-testing from the fourth generation onwards and ending with a few promising breeding clones after the 7^th^ year of field-testing and selection to include them in multi-site trials in the target population of environments. The genomic prediction accuracy over the two years within each site suggests that it will be possible to select (based on GEBV models) in early generation trials for each target population of environments. Furthermore, as per previous GP accuracy estimates (Ortiz et al., 2022; Selga et al., 2022) and these results, it seems that GEBV for selection will be useful from T_3_ onwards rather than in T_1_ or even in T_2_. Hence, as shown herein, genomic selection appears to be feasible in potato breeding when using elite bred germplasm.

## 5. CONCLUSION

We investigated the accuracy of four genome-based prediction models including either Hadamard or Kronecker product matrices for assessing GE. Several prediction problems were analyzed for the GP accuracy of each of the four models. We investigated the prediction set of locations in year 2021 from locations in year 2020 using the four GP models combined with two prediction sets (100% and 70%) using both ST and MT. The ST model *M3Sp* was the best genomic predicted, followed by *M1Sp* and*M1Sa* at HEL in 2021. In terms of MT GP accuracy, *M4Mp\** was the best for weight of tubers with 40−50mm, above 60 mm and total tuber weight irrespective of size, and very similar to tuber flesh starch. The GP accuracy of the seven traits at MOS in 2021 indicated that the best models for predicting the majority of the seven traits were ST *M3Sp* and *M2Sp*, except for weight of tubers with 50−60mm, above 60mm, and total tuber weight, where MT model *M4Mp\** was the best GP model. The traits with higher phenotypic correlations between location UM 2021 and those at HEL, MOS, and UM in 2020 are weight of tubers with 50−60 mm, below 40 mm, and above 60 mm. The best model-method for predicting the majority of the seven traits was MT *M4Mp\** because it allows exchange information between traits and environments followed by *M3Sp* and *M2Sp* that efficiently used information between environments. According with Cuevas et al (2017) it was expected *M3Sp* producing better or similar GP accuracy than *M2Sp*.

## 6. DATA AVAILABILITY STATEMENT

DNA marker and phenotypic data for each year within sites are stored at https://hdl.handle.net/11529/10548784

## 9. AUTHOR CONTRIBUTIONS

RO conceptualized the research, and together with JCr and FR did the experimental designs for all trials. FR and RO carried out evaluations and data recording. JCu, JCr and RO did the analysis and interpretation of the research results. All authors wrote the manuscript under the leadership of JCr.

